# SF3b inhibitors suppress proliferation of non-small cell lung cancer cells irrespective of EGFR mutation status

**DOI:** 10.64898/2026.10.29.762206

**Authors:** Wataru Fukagawa, Yuta Horita, Hiroyoshi Harada, Masayoshi Mori, Yusuke Murata, Kenji Ohe

## Abstract

Epidermal growth factor receptor tyrosine kinase inhibitors (EGFR-TKIs) are standard care for EGFR-mutant non-small cell lung cancer (NSCLC), but acquired resistance limits their durability, and alternatives that do not depend on EGFR kinase activity are needed. We asked whether inhibitors of SF3b, a core component of the U2 small nuclear ribonucleoprotein particle, act on NSCLC cells regardless of EGFR genotype. Three lung adenocarcinoma lines were treated with pladienolide B or herboxidiene at 0.01, 0.1 and 1 µM: PC9 and HCC827, both carrying an EGFR exon 19 deletion and both inhibited by gefitinib at nanomolar concentrations, and A549, which is EGFR wild-type and gefitinib-insensitive. Proliferation was followed for three days by crystal violet staining, EGFR intron 2 retention was assayed by RT-PCR, and EGFR protein was measured by western blotting. Both compounds produced a concentration-dependent increase in the intron 2-retained product in all three lines, and both held proliferation at or near the day-0 cell number in all three, at concentrations down to 0.01 µM. EGFR protein was lost in PC9, reduced in A549, and only partly reduced in HCC827, where a residual band persisted in lanes whose growth was fully suppressed. Since A549 is EGFR wild-type, and published knockdown data show that removing EGFR does not suppress proliferation of wild-type lines, the growth inhibition cannot be attributed to loss of EGFR alone. We read EGFR intron 2 retention as a marker of target engagement rather than as the mechanism of the growth effect. To place that read-out in context without further experiments, we reanalysed public RNA-seq data from Rh18 cells treated with the spliceostatin analogue sudemycin D1 and measured all 27 EGFR introns. EGFR as a gene was not unusually sensitive to SF3b inhibition (Mann-Whitney p = 0.309 against 2,041 genome-wide introns), and intron 2 was among the least responsive junctions within it, while introns 1, 3, 11, 13, 14 and 19 each rose by more than 15 percentage points. Intron 2 carries an unusually strong 3′ splice site (88.2nd percentile genome-wide), which is consistent with that resistance. Intron 2 therefore reports engagement in our hands but should not be assumed to be the most sensitive EGFR reporter available. These are the results of single experiments without replicates or statistical testing and should be read as preliminary.

## Introduction

Lung cancer remains the leading cause of cancer death worldwide, and non-small cell lung cancer (NSCLC) accounts for the large majority of cases. In lung adenocarcinoma, activating mutations in the epidermal growth factor receptor (EGFR) gene — most commonly exon 19 deletions and the L858R substitution — define a subgroup for which EGFR tyrosine kinase inhibitors (EGFR-TKIs) are first-line therapy. Gefitinib, erlotinib and afatinib were followed by osimertinib, which is active against the T790M gatekeeper mutation that accounts for roughly half of acquired resistance to earlier-generation agents.

Resistance nevertheless emerges in essentially all patients. Beyond T790M, it arises through amplification of MET, activation of FGFR and other bypass receptors, mutations in downstream effectors such as KRAS, NRAS and BRAF, and histological transformation. Each mechanism restores signalling downstream of, or parallel to, the receptor, so a therapeutic strategy that does not depend on inhibiting EGFR kinase activity would be useful whatever the resistance mechanism.

Pre-mRNA splicing offers one such strategy. Splicing factor 3b (SF3b) is a multi-subunit component of the U2 small nuclear ribonucleoprotein particle that recognises the branch point during spliceosome assembly. Somatic mutations in its largest subunit, SF3B1, were identified in myelodysplastic syndrome with ring sideroblasts and are now recognised across several haematological and solid tumours, which established the spliceosome as a druggable target. Natural products that bind SF3b — pladienolide B, herboxidiene and their derivatives — shift U2 snRNP away from the canonical branch point, and the consequence is not uniform suppression of splicing but selective alteration of individual introns, some of which are retained while others are spliced normally.

We had previously observed, in unpublished work, that SF3b inhibitors cause retention of EGFR intron 2 in SH-SY5Y cells, confirmed by sequencing. Retention of intron 2 introduces a premature termination codon, which predicts degradation of the transcript by nonsense-mediated decay and a corresponding loss of EGFR protein. If this occurred in NSCLC cells, SF3b inhibition would remove the receptor itself rather than inhibit its kinase, and would in principle be unaffected by kinase-domain resistance mutations.

We therefore treated three lung adenocarcinoma cell lines of differing EGFR genotype and EGFR-TKI sensitivity — PC9 and HCC827, both EGFR exon 19 deletion and both highly gefitinib-sensitive, and A549, EGFR wild-type and gefitinib-insensitive — with pladienolide B and herboxidiene, and asked whether EGFR intron 2 retention, loss of EGFR protein and growth inhibition occurred together.

## Materials and methods

### Compounds

Pladienolide B (Cayman Chemical) and herboxidiene (FOCUS Biomolecules) were dissolved in DMSO. DMSO alone served as the vehicle control.

### Cell lines

Three human lung adenocarcinoma lines were used. All were maintained in DMEM (Nacalai Tesque) with 10% FBS (Japan Bio Serum) at 37 °C under 5% CO_2_.

| **Cell line** | **EGFR** | **Other** | **EGFR-TKI sensitivity** |
|---|---|---|---|
| PC9 | exon 19 deletion, p.Glu746_Ala750del | — | sensitive |
| HCC827 | exon 19 deletion, p.Glu746_Ala750del | TP53 p.Val218del | sensitive |
| A549 | wild-type | KRAS p.Gly12Ser | insensitive |

### Proliferation assay

Cells were seeded in 24-well plates at 2 × 10^4^ cells per well and treated with pladienolide B or herboxidiene at 0.01, 0.1 or 1 µM, or with DMSO, for up to 72 h. Viable cells were stained at 24, 48 and 72 h with a Crystal Violet Assay Kit (Abcam) and absorbance was read at 595 nm on a microplate reader (Tecan). Values are expressed as the ratio to the 0 h reading of the same condition.

Growth inhibition at each concentration was calculated from the day-3 reading (day 2 for the herboxidiene series in HCC827) as 1 − (S_h_ − S_0_) / (S_v_ − S_0_), where S_h_ is the treated value, S_v_ the vehicle value at the same time point and S_0_ the 0 h value of the same series. IC50 was taken as the concentration giving 50% inhibition, interpolated linearly between the two bracketing concentrations on a log scale. Where 50% inhibition was already exceeded at the lowest concentration tested, the IC50 is reported as below that concentration rather than extrapolated.

### RT-PCR

Cells were treated for 24 h, lysed in TRIzol (Invitrogen) and extracted with chloroform. Total RNA was purified from the aqueous phase and 1 µg was reverse transcribed with ReverTra Ace (Toyobo). PCR was performed with Ex Taq (Takara) for 30 cycles, with annealing at 60 °C and extension for 60 s, using the primers below; products were resolved on an agarose gel.

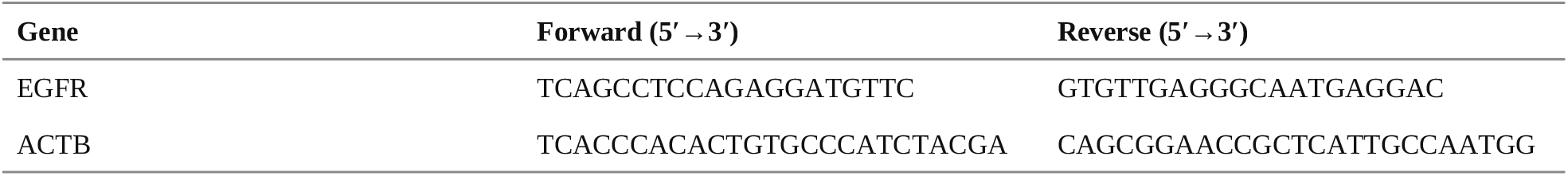

The EGFR primers lie in exons 2 and 3 and span intron 2. In silico PCR against the human genome (hg38) returns a single genomic product of 1006 bp at chr7:55,142,343-55,143,348, made up of 95 bp of exon 2, intron 2 of 867 bp with canonical GT and AG termini, and 44 bp of exon 3. The spliced product is therefore 139 bp and the fully intron 2-retained product 1006 bp, and the exonic sequence reads in a single open frame across the junction (… SFLK | TIQEVAGY…), confirming that the two primers sit in consecutive exons.

### Western blotting

Protein was recovered by acetone precipitation from the interphase of the same TRIzol extraction, dissolved in urea sample buffer (10 mM Tris-HCl pH 7.5, 25 mM KCl, 5 mM MgCl_2_, 6 M urea) and sonicated to shear co-precipitated DNA and RNA. Protein was quantified with the Pierce 660 nm Protein Assay (Thermo Fisher). Approximately 20 µg per lane was separated by SDS-PAGE and transferred to PVDF by tank transfer. Membranes were blocked with Blocking One (Nacalai Tesque) for 10 min at 4 °C, incubated with primary antibody (1:1,000) for 16 h at 4 °C and with secondary antibody (1:5,000) for 1 h at room temperature, and developed with Chemi-Lumi One L (Nacalai Tesque). Antibodies: anti-EGFR (4267T, Cell Signaling Technology), anti-ACTB (A00702, CGCS), anti-mouse IgG HRP-linked (7076S, Cell Signaling Technology).

### Replication

Every experiment reported here was performed once. The proliferation curves, the RT-PCR panels and the western blots are each a single run, presented as a representative observation without densitometric quantification or statistical testing. Independent replication has not been established for any of the results in this report.

### Reanalysis of public RNA-seq data

Raw RNA-seq reads for GSE102539 (Rh18 rhabdomyosarcoma cells treated with DMSO or 1 µM sudemycin D1 for 8 h, in triplicate) were obtained from the Sequence Read Archive (runs SRR5927887-SRR5927892) and sample-to-group assignment was read from the GEO record. Retention was quantified without alignment. For each intron a spliced 30-mer (15 nt of upstream exon joined to 15 nt of downstream exon) and a retained 30-mer (the same upstream 15 nt joined to the first 15 nt of the intron) were generated together with their reverse complements, and reads were matched against the pattern set by exact string search, counting both strands. Retention was expressed as the percent intron retention, PIR = IR / (IR + spliced) x 100, and drug effect as ΔPIR, the treated minus control difference in pooled replicates. All 27 introns of EGFR transcript NM_005228.5 were measured. As a reference distribution we used a panel of 6,000 introns sampled across the range of published responses to the SF3b inhibitor E7107, of which 2,041 reached the coverage threshold of 15 junction reads in both arms in these samples. Introns with fewer than 15 reads in either arm were excluded. Exact bounds on a zero count used the Clopper-Pearson one-sided 95% interval. Splice sites were scored with position weight matrices for the 9-nt donor and 23-nt acceptor built from 703 annotated human introns, and EGFR scores were expressed as percentiles of the 6,000-intron panel. Group comparisons used the Mann-Whitney U test.

## Results

In all three lines the RT-PCR across intron 2 resolved three products: the 139 bp spliced product, the 1006 bp intron 2-retained product, and an intermediate band. The intermediate band was sequenced and reported as a non-specific product. Since the amplicon contains only two exons, it cannot represent an exon-skipped form of this transcript.

### PC9 cells

DMSO-treated PC9 cultures grew about 4.7-fold over three days (Figure 1A). Pladienolide B held proliferation at the day-0 level at every concentration tested, including the lowest, 0.01 µM (Figure 1A). Herboxidiene did the same at 0.1 and 1 µM, but at 0.01 µM growth was slowed rather than abolished, reaching about 2.6-fold by day 3 (Figure 1B). Because 0.01 µM pladienolide B was already maximally effective, the concentration range did not capture the lower end of the dose-response for that compound.

**Figure 1.**
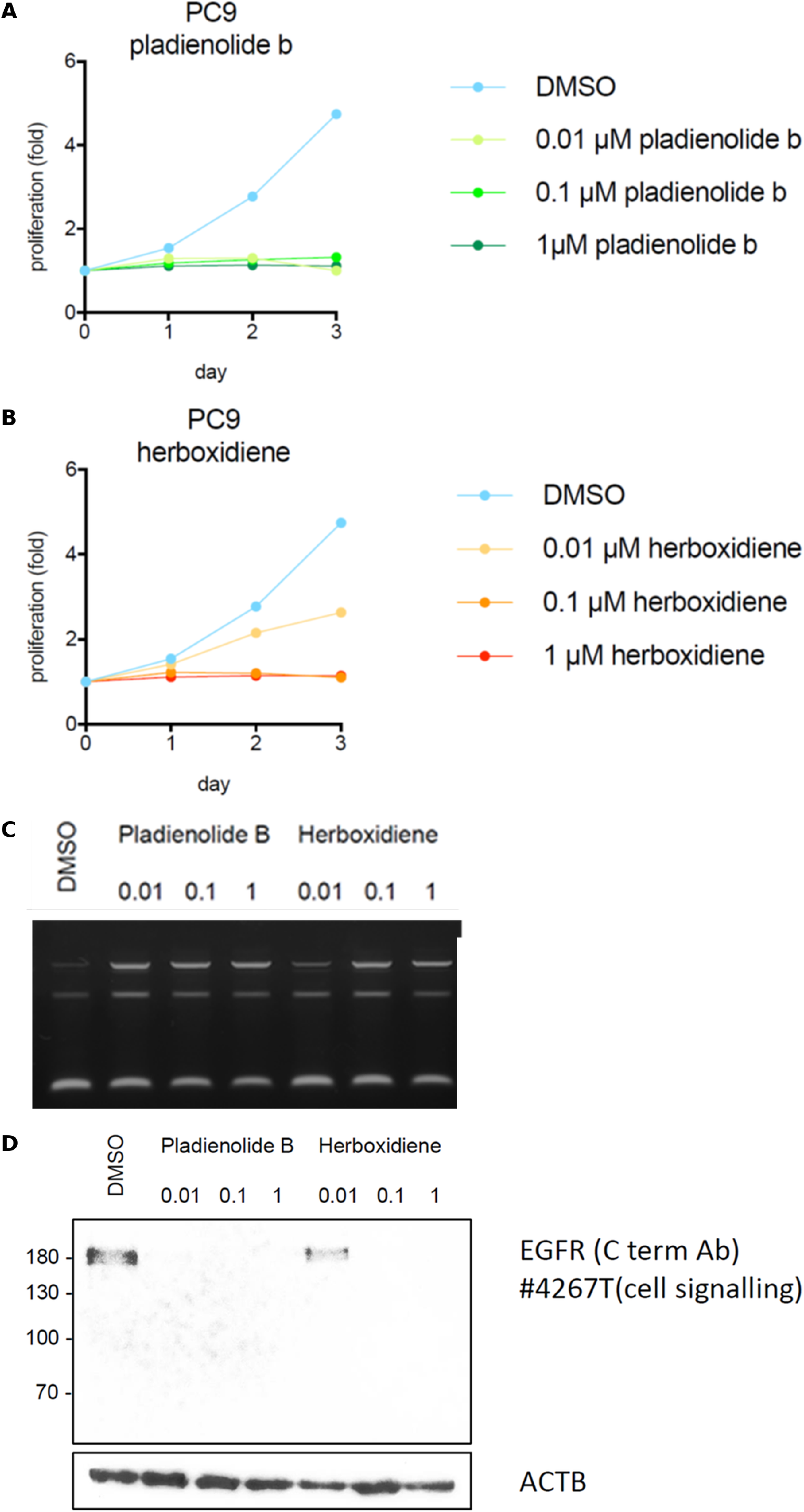
SF3b inhibitors in PC9 cells (EGFR exon 19 deletion, gefitinib-sensitive). (A) Proliferation over three days with pladienolide B at 0.01, 0.1 and 1 µM against DMSO, measured by crystal violet staining at 595 nm and expressed as the ratio to the 0 h reading. (B) The same for herboxidiene. (C) RT-PCR across EGFR intron 2, with primers in exons 2 and 3, after 24 h of treatment. The slowest band is the intron 2-retained product (1006 bp) and the fastest is the spliced product (139 bp); the intermediate band was sequenced and reported as non-specific. (D) Western blot for EGFR (C-terminal antibody 4267T) and ACTB after 24 h of treatment, with molecular mass markers in kDa. Single experiment; no replicates and no statistical testing.

The intron 2-retained product increased with concentration for both compounds, but was weaker at 0.01 µM herboxidiene than at the higher concentrations (Figure 1C). EGFR protein was lost at all three concentrations of pladienolide B and at 0.1 and 1 µM herboxidiene, and persisted at 0.01 µM herboxidiene (Figure 1D); ACTB was unchanged. The one condition in which EGFR protein remained is the same one in which intron 2 retention was weakest and growth was not fully suppressed.

### HCC827 cells

HCC827 cells carry the same EGFR exon 19 deletion as PC9. DMSO-treated cultures grew about 2.2-fold over three days (Figure 2A). Pladienolide B held proliferation at or just below the day-0 level at all three concentrations (Figure 2A). Herboxidiene did the same at 0.1 and 1 µM; at 0.01 µM a small increase to about 1.15-fold was seen by day 2 (Figure 2B). The day-3 samples for the herboxidiene series were not recovered, so that panel ends at day 2.

**Figure 2.**
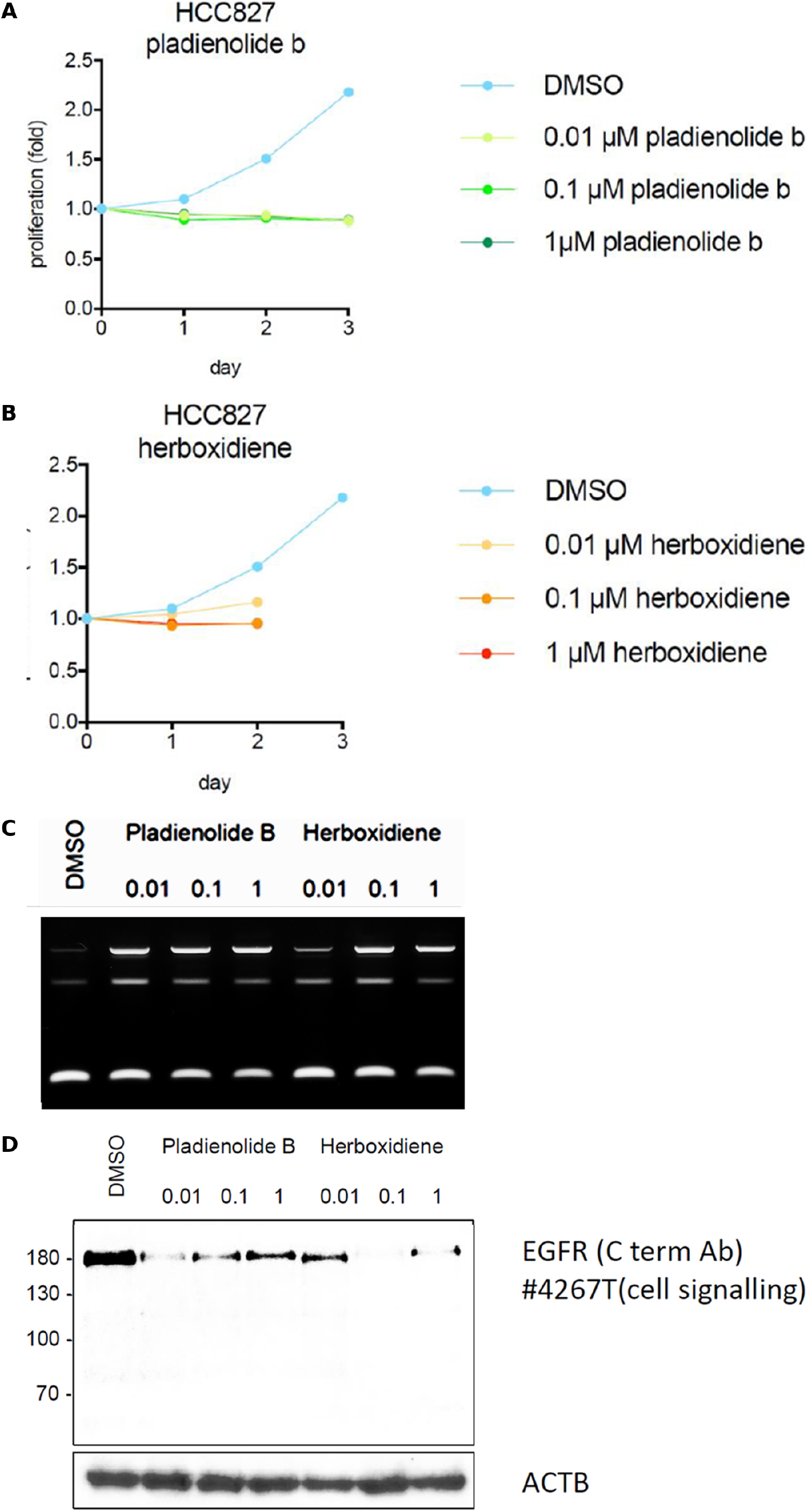
SF3b inhibitors in HCC827 cells (EGFR exon 19 deletion, TP53 deletion, gefitinib-sensitive). Panels as in Figure 1. The day-3 samples of the herboxidiene series in (B) were not recovered.

The intron 2-retained product increased with concentration for both compounds, again more weakly at 0.01 µM herboxidiene, matching PC9 (Figure 2C). EGFR protein, in contrast to PC9, was not abolished: a residual band was present in most treated lanes, strongest at 0.1 and 1 µM pladienolide B (Figure 2D). ACTB was unchanged. Growth was nonetheless suppressed in those same lanes.

### A549 cells

A549 cells are EGFR wild-type and gefitinib-insensitive. DMSO-treated cultures grew about 2.5-fold over three days (Figure 3A). Both compounds suppressed growth at all three concentrations: from day 1 onward the treated cultures stayed at or below the day-0 value, with a slight upturn by day 3 at the lower concentrations (Figure 3A, 3B).

**Figure 3.**
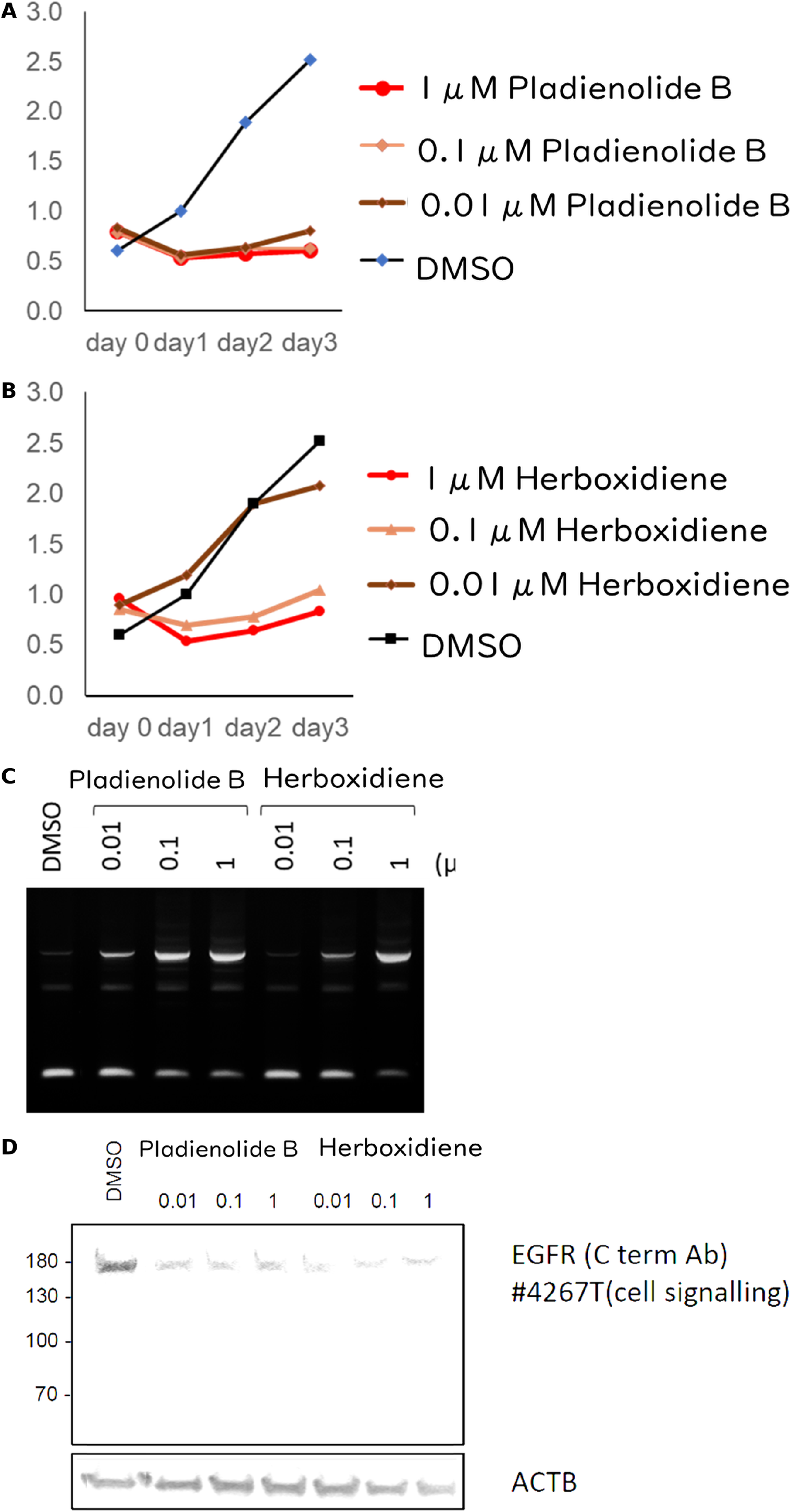
SF3b inhibitors in A549 cells (EGFR wild-type, gefitinib-insensitive). Panels as in Figure 1.

The intron 2-retained product increased with concentration for both compounds, and the gradient was the steepest of the three lines (Figure 3C). Unlike PC9 and HCC827, the weakest retention here was at 0.01 µM pladienolide B rather than at 0.01 µM herboxidiene. EGFR protein was reduced in all treated lanes relative to DMSO, to an extent comparable with PC9 (Figure 3D).

### Summary across the three lines

#### Potency

IC50 values calculated from the proliferation curves are given below. In five of the six compound-line combinations, 50% inhibition was already exceeded at 0.01 µM, the lowest concentration tested, so the IC50 is bounded rather than determined; only herboxidiene in A549 gave a curve that crossed 50% within the range tested. The design therefore does not define the potency of either compound, and pladienolide B in particular needs to be examined below 0.01 µM.

| **Cell line** | **Compound** | **Inhibition at 0.01 µM** | **Inhibition at 0.1 µM** | **Inhibition at 1 µM** | **IC50** |
|---|---|---|---|---|---|
| PC9 | pladienolide B | 99% | 93% | 92% | < 0.01 µM |
| PC9 | herboxidiene | 57% | 96% | 97% | < 0.01 µM |
| HCC827 | pladienolide B | 110% | 109% | 110% | < 0.01 µM |
| HCC827 | herboxidiene | 71% | 112% | 112% | < 0.01 µM |
| A549 | pladienolide B | 99% | 110% | 109% | < 0.01 µM |
| A549 | herboxidiene | 39% | 91% | 106% | 0.016 µM |

Inhibition above 100% means the treated culture fell below its own 0 h reading. HCC827 herboxidiene values are from day 2, the last time point recovered for that series; all others are from day 3.

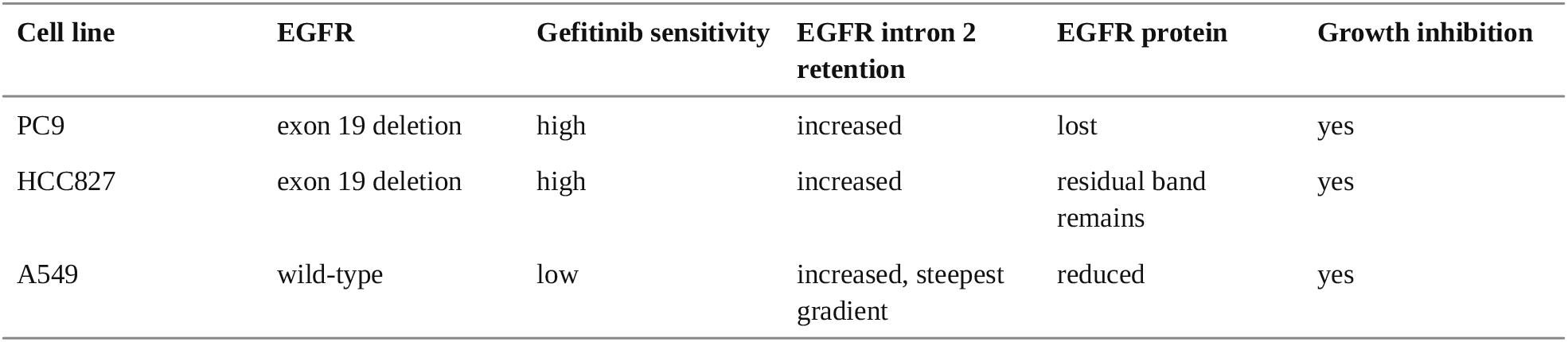

### EGFR intron 2 compared with the rest of the gene

The intron 2 primer pair was chosen because it was already in hand, not because intron 2 was known to be the most drug-responsive EGFR intron. To ask how it compares with the other 26, we reanalysed public RNA-seq data rather than perform further experiments (Fig. 4). We used GSE102539, in which Rh18 rhabdomyosarcoma cells were treated with the spliceostatin analogue sudemycin D1 against a vehicle control from the same laboratory; sudemycin D1 is a different chemotype from pladienolide B and herboxidiene but engages the same branch-point pocket of SF3b.

**Figure 4.**
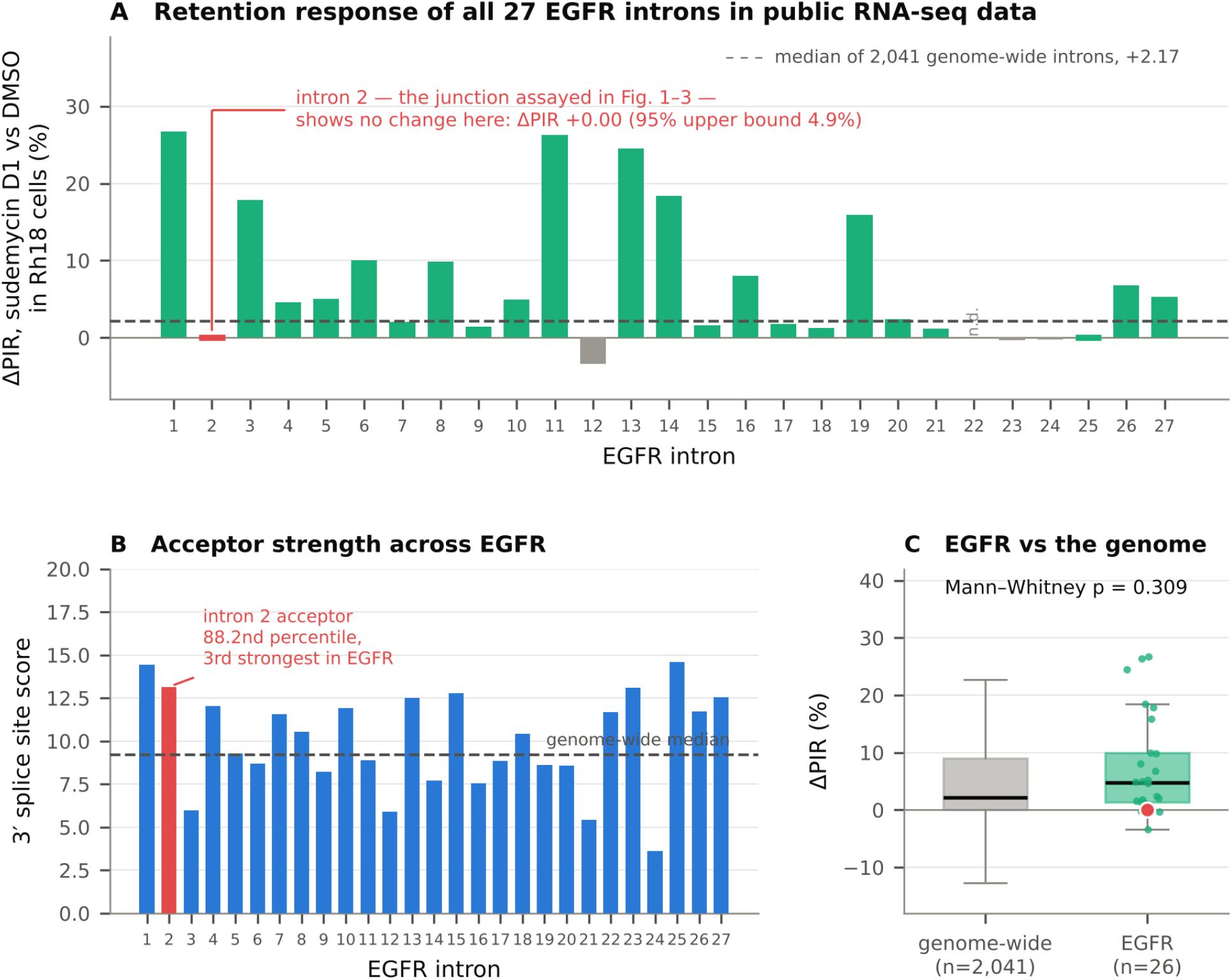
EGFR intron 2 compared with the rest of the gene in public RNA-seq data. Reanalysis of GSE102539 (Rh18 rhabdomyosarcoma cells, 1 µM sudemycin D1 or DMSO for 8 h, triplicate). (A) Change in percent intron retention (ΔPIR) for each of the 27 EGFR introns. Intron 2, the junction assayed in Figures 1C, 2C and 3C, is marked in red; its value is a measured zero and is drawn as a flat marker to distinguish it from a missing value. n.d., not determined for lack of coverage. Dashed line, median of the 2,041-intron reference panel in the same samples. (B) 3′ splice site score for each EGFR intron, with intron 2 marked; dashed line, median of the reference panel. (C) Distribution of ΔPIR for EGFR introns against the reference panel; red point, intron 2.

EGFR responded to the drug, but unremarkably. Its median ΔPIR across the 26 measurable introns was +4.76 against +2.17 for the 2,041-intron reference panel in the same samples, placing EGFR at the 63.2nd percentile; the difference was not significant (Mann-Whitney p = 0.309; Fig. 4C). By this measure EGFR is a modestly above-median gene rather than a privileged target of SF3b inhibition.

Within EGFR the response was very uneven (Fig. 4A). Introns 1 (+26.71), 11 (+26.33), 13 (+24.50), 14 (+18.42), 3 (+17.87) and 19 (+15.90) all rose substantially, whereas intron 2 showed no detectable change: ΔPIR +0.00, ranking 22nd of 26, with zero intron-retaining reads among 256 junction reads in vehicle and among 60 under drug. The exact one-sided 95% upper bounds on PIR are 1.2% and 4.9% respectively, so a response of the size seen at introns 1 or 11 is excluded in these cells, although a small one is not.

One sequence feature is consistent with that resistance. Drug-sensitive introns have been reported to carry weak 3′ splice sites, and the intron 2 acceptor is unusually strong: its score places it at the 88.2nd percentile of the genome-wide panel and third of the 27 EGFR introns (Fig. 4B). Its length cuts the other way, since 867 nt is short and short introns have been reported as more sensitive, so the determinants account for the result only in part.

This analysis uses a different cell line, a different compound and a different assay from Figures 1 to 3, and does not contradict the retention we observed there. It does indicate that intron 2 is unlikely to be the most sensitive EGFR reporter of SF3b engagement.

## Discussion

Pladienolide B and herboxidiene suppressed proliferation in all three lung adenocarcinoma lines, and the effect did not follow EGFR genotype. PC9 and HCC827 carry an activating exon 19 deletion and are inhibited by gefitinib at nanomolar concentrations; A549 is EGFR wild-type and requires micromolar gefitinib. All three were held at or near their day-0 cell number by 0.01 µM pladienolide B. Whatever the SF3b inhibitors are doing, it is not confined to cells whose growth depends on mutant EGFR.

Both compounds induced EGFR intron 2 retention in all three lines, and the amount of retained product rose with concentration. Intron 2 retention introduces a premature termination codon, which predicts degradation of the transcript by nonsense-mediated decay and a fall in EGFR protein. That fall was seen in PC9, where the band was lost outright, and in A549, where it was clearly reduced. In PC9 the three readouts moved together: at 0.01 µM herboxidiene, the one condition with weak intron retention, EGFR protein persisted and growth was only slowed. That internal consistency is the strongest evidence here that the compounds engage the EGFR transcript.

Two observations argue that this is not the whole mechanism. First, in HCC827 a residual EGFR band remained in most treated lanes, yet proliferation was suppressed as completely as in PC9. Second, and more decisively, A549 is EGFR wild-type and gefitinib-insensitive. Shien and colleagues silenced EGFR in 13 NSCLC lines and found that viability fell in six of eight EGFR-mutant lines but in none of the wild-type lines except NCI-H411, in which EGFR is constitutively phosphorylated. Removing EGFR from a line such as A549 would not be expected to stop it growing, so the growth inhibition seen here in A549 cannot be attributed to the loss of EGFR protein even though that loss occurred.

The most economical reading is that EGFR intron 2 retention is a reliable marker of target engagement, and in EGFR-dependent cells may well contribute to the growth effect, but that it is not sufficient to explain the effect across genotypes. SF3b inhibitors alter branch-point selection globally, and the transcripts that respond differ between introns and between cell types; many genes required for proliferation are likely affected at once. Identifying which of them matters would require a transcriptome-wide comparison of treated and untreated cells in each line, which is the obvious next step. A second step would be to test whether the growth effect survives in lines carrying T790M or MET amplification, since a strategy that does not depend on EGFR kinase activity is only interesting if it holds where the TKIs fail. It would also be worth asking whether other introns of EGFR, and the introns of the other actionable drivers in lung adenocarcinoma — ALK, ROS1, BRAF, MET, KRAS, NTRK — respond in the same way.

Several limitations should be weighed against these observations. Each assay was performed once per line, with no replicates and no statistical testing, so the magnitude of every effect reported here is provisional and the smaller differences between concentrations should not be read as real. Because 0.01 µM pladienolide B was already maximally effective in all three lines, the experiment does not define a lower bound for its activity, and the picomolar range should be examined. The identity of the intron 2-retained product was established previously in SH-SY5Y cells by sequencing but was not re-sequenced in the lines used here, and the product was not shown to be degraded by nonsense-mediated decay, which would require a decay-inhibition control. The EGFR antibody recognises the C-terminus only; a truncated product initiated downstream would not be detected. Phosphorylated EGFR was not measured, so the functional state of the receptor remaining in HCC827 is unknown. The western blots were not quantified by densitometry, which is the first thing to add. No non-transformed cell line was included, so nothing here speaks to a therapeutic window, and no EGFR-independent lung line was run as a negative control.

The public-data analysis qualifies our read-out rather than our result. We observed a concentration-dependent increase in the intron 2-retained product in all three lung lines with two different compounds (Figs. 1C, 2C, 3C), so that junction plainly can report engagement in those settings. What the reanalysis adds is that in a different cell line and with a third compound, intron 2 was among the least responsive EGFR introns while six others rose by more than 15 percentage points, so an assay reading two or three EGFR junctions would have more headroom than intron 2 alone; introns 11 and 1 would be the obvious additions. The reanalysis also shows that EGFR as a gene is not unusually sensitive to SF3b inhibition. That is consistent with, rather than contrary to, our main conclusion: if EGFR were a privileged target of these compounds, the uniform growth inhibition across genotypes would be harder to reconcile with the published finding that removing EGFR does not suppress proliferation of wild-type lines.

SF3b-targeting agents have reached clinical study. E7107, a pladienolide B derivative, entered a phase I trial that was stopped after visual disturbance in treated patients, and the oral SF3B1 modulator H3B-8800 completed a first-in-human study in myeloid neoplasms without meeting its efficacy endpoint, though some patients became transfusion-independent. Splicing is essential in every cell, so the therapeutic window for any such agent is the central question, and it is not addressed by the experiments reported here. We post this work as a preprint because, as far as we can establish, SF3b inhibitors have not previously been tested in lung cancer cell lines, and because the observation that the effect is indifferent to EGFR genotype seems worth recording while those questions remain open.

## Supporting information

Supplementary data and code for Figure 4

## Declarations Author contributions

Conceptualization: K.O. Methodology: K.O. Investigation: W.F. and K.O. Formal analysis: K.O. Visualization: K.O. Writing — original draft: K.O. Supervision and project administration: K.O. Y.H., H.H., M.M. and Y.M. advised on study design and interpretation. Writing — review and editing: all authors. All authors read and approved the final manuscript.

## Data availability

All data generated in this study are presented in the figures. The public data reanalysed in Figure 4 are available from the Gene Expression Omnibus under GSE102539 and from the Sequence Read Archive under runs SRR5927887-SRR5927892; the per-intron values plotted in Figure 4 and the scripts used to produce them are provided as supplementary files. The original gel and blot images are available from the corresponding author on request.

