## Supplementary figures and images for "SF3b inhibitors suppress proliferation of non-small cell lung cancer cells irrespective of EGFR mutation status"

### Figure4_EGFR_public.png

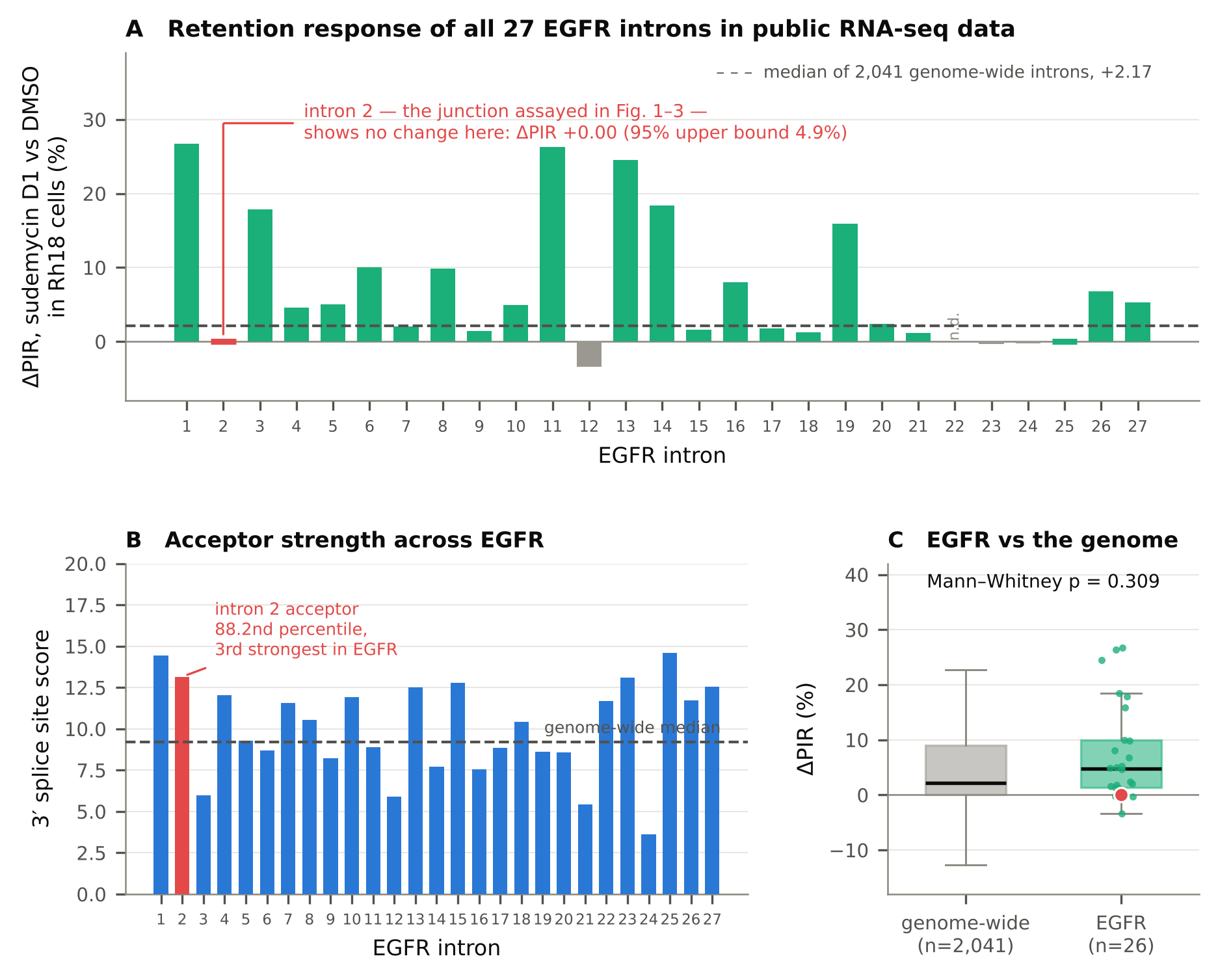
